# Energy savings across the cortex for confidently predicted visual input

**DOI:** 10.1101/2023.12.08.570804

**Authors:** André Hechler, Floris P. de Lange, Valentin Riedl

## Abstract

Our expectations about the world influence how we interpret visual information, improving the speed and accuracy of perception. However, the underlying neural activity requires energy which is strictly limited in the brain. While predictive processing is a prevalent framework to explain perception, it remains unclear whether it also serves energy-efficient processing.

Here, we employed metabolic brain imaging to quantify oxygen consumption during visual perception under varying levels of input predictability and subjective uncertainty. For three days, we presented participants with object sequences that were either predictable or unpredictable, and assessed their performance and confidence in predicting follow-up objects from partial sequences. On the fourth day, we first tested for behavioral consequences of predictability. We found that subjects detected predictable objects quicker than unpredictable ones. We then quantified cortical oxygen consumption during passive viewing of predictable, unpredictable or surprising sequences. Despite highly similar sensory load, predictable visual input elicited reduced oxygen metabolism when subjects were confident, across both sensory and higher cognitive areas. Crucially, this summed up to cortical energy savings of up to 12%, or 118 μmol oxygen per minute, given average brain size. In contrast, cost increases due to surprising input were restricted to a network of fronto-parietal areas.

In summary, we found that predictive processing enhances behavioral performance and notably reduces signaling costs, moderated by subjective confidence. This suggests that examining energy efficiency alongside behavioral performance may uncover novel computational strategies of human cognition and behavior.

## Introduction

Understanding how our environment changes over time is a crucial ability for survival, allowing us to anticipate the future and adapt our behavior accordingly. This necessitates the extraction of patterns or rules from our sensory input, leading to the expectation that certain observations tend to follow each other in time. Research on visual perception showed that humans are skilled at learning probabilistic and deterministic patterns (Fiser & Aslin, 2002; Maheu et al., 2022; R. Wang et al., 2017) This impacts both behavior and perception itself: Expected stimuli are detected quicker and with higher accuracy and ambivalent percepts are biased towards prior beliefs (de Lange et al., 2018; Turk-Browne et al., 2005).

Over the last decades, evidence converged that we maintain a *generative model* of the world, an internal representation of the external processes we perceive (Dayan et al., 1995; Friston, 2010). This model functions according to Bayesian updating, integrating prior knowledge and current evidence to provide an optimal estimate (Knill & Pouget, 2004; Rescorla, 2021). While a dedicated framework for the neurobiological implementation of this process exists (*predictive coding*; Bastos et al., 2012; Friston, 2005; Rao & Ballard, 1999), a crucial factor of its validity remains underexplored: Does predictive processing primarily serve behavioral performance, potentially at higher signaling cost, or is it instead an efficient framework that enables the brain to conserve its limited energy resources?

The brain, in relation to its size, is the most energetically demanding organ in the human body, relying on a constant supply of energy substrates. It consumes 20% of the body’s energy intake (Rolfe & Brown, 1997), and about 75% of this energy is devoted to neuronal activity (Howarth et al., 2012). Healthy brain function relies heavily on the steady provision of glucose and oxygen, the essential components of ATP synthesis (Shobatake et al., 2022; Warren & Frier, 2005). As a result, reliable energy availability has played a crucial role in the evolution of brain anatomy and cognitive function (Niven & Laughlin, 2008; Quintela-López et al., 2022).

Theoretical work suggests that predictive coding promotes energy-efficient perception (Chalk et al., 2018; Manookin & Rieke, 2023; Sengupta et al., 2013). While expected neural signals are suppressed, feed-forward transmission of input that deviates from expectations is prioritized. Consistent with this theory, studies using neuroimaging and single cell recordings indicate that a given stimulus elicits a weaker response when it is expected compared to when it is surprising (Alink et al., 2010; Kaposvari et al., 2018; Meyer & Olson, 2011; Richter et al., 2018). However, inferring the energy metabolic effects from this evidence is complicated, especially when considering processing across the cortical hierarchy.

To thoroughly assess the energy efficiency of predictive processing, it is essential to measure the absolute metabolic rates throughout the entire cortex. This has been challenging using current methods for studying predictive processing. Firstly, human studies mainly relied on standard fMRI and its BOLD (blood-oxygenation level dependent) signal (Alink et al., 2010; Richter et al., 2018). Crucially, the BOLD signal cannot disentangle oxygen consumption from confounding effects of oxygen supply and blood flow (Drew, 2019, 2022). Furthermore, variations in vasculature and hemodynamic response make it impossible to compare BOLD signal amplitudes across different brain regions or individuals (Ances et al., 2008; Devonshire et al., 2012). Therefore, BOLD differences during predictive processing reflect general changes in regional oxygenation, not energy metabolism. Secondly, invasive recordings in animals directly capture neuronal activity, but they are limited to individual cells or small populations (Kaposvari et al., 2018; Meyer & Olson, 2011). Predictive coding is a multi-level process (Friston, 2005; Rao, 2024), and it is yet unclear how local effects scale when considering activity beyond sensory areas. Consequently, a quantitative approach to measure metabolic rates of energy substrates across the entire brain is necessary to investigate the system-wide energy efficiency of predictive processing.

In the present study, we employed an advanced metabolic imaging method to measure the cortical energy consumption during visual stimulation of varying predictability. We used multiparametric quantitative BOLD (mqBOLD) which quantifies the consumption of oxygen across the brain (Christen et al., 2012; Hirsch et al., 2014; Kaczmarz et al., 2020). As at least 90% of the brain’s energy metabolism is oxidative (Blazey et al., 2018; Lin et al., 2010), mqBOLD provides direct access to the brain’s energy turnover. To manipulate the predictability of the input, we extended an established statistical learning paradigm (Meyer & Olson, 2011; Richter et al., 2018), in which participants are presented with visual object sequences. Over three days, participants learned the underlying patterns and reported their subjective confidence when predicting the next object in a sequence. We used these ratings as a measure of interindividual differences in prediction ability (Fleming, 2024). Confidence ratings are reflective of perceived uncertainty and the subjective probability of being correct (Navajas et al., 2017; Pouget et al., 2016). Furthermore, they provide information about the relative weighting of prior knowledge and evidence (Meyniel, Sigman, et al., 2015). To study the processing efficiency of various levels of predictability, we also presented sequences with random order and sequences that deviated from the learned patterns.

We found that the energetic cost of predictive processing depends on the level of subjective confidence in predicting the presented patterns. With high confidence, energy consumption during predictable input decreased by up to 12% when summed across the cortex. At the same time, participants also displayed behavioral benefits of predictive structure, similarly improving with confidence. These results show that predictive processing optimizes both performance and energetic efficiency, in line with the evolutionary pressure for sensory systems to be accurate without prohibitively high cost.

## Results

### Confidence and processing speed increase with repeated exposure to visual sequences

During the first part of the study, participants completed a three-day sequence learning phase, which was conducted online (Methods). We presented visual sequences of everyday objects that either followed a deterministic pattern (predictable condition, P) or a random pattern (unpredictable condition, U) (Figures 1a and b). To ensure vigilance and gaze fixation, participants had to react to occasionally presented (10 % of trials) upside-down objects, which occurred irrespective of sequence predictability. These *catch trials* were detected with high accuracy (mean: 96.3%, SD: 0.02), indicating that participants stayed concentrated during the online training. Neither accuracy nor reaction time differed between conditions (reaction time: t(*40*)=-1.4, p=.17; accuracy: t(*40*)=1.51, p=.14). We assessed the trajectory of sequence learning on a daily basis, both on an objective and a subjective scale. To this end, we presented participants with incomplete sequences and prompted them to choose the correct follow-up object. For each choice, we assessed confidence ratings as an indicator of uncertainty regarding knowledge of the underlying patterns (Figure 1b). Over the training days, confidence ratings increased for predictable visual sequences and remained constant for unpredictable ones (Figure 1d). We also saw an objective improvement in pattern completion (Figure 1e) which was highly correlated with confidence (r=0.75, p<0.001). Since we did not provide any feedback, this shows that participants were accurate in assessing their performance.

**Fig 1.**
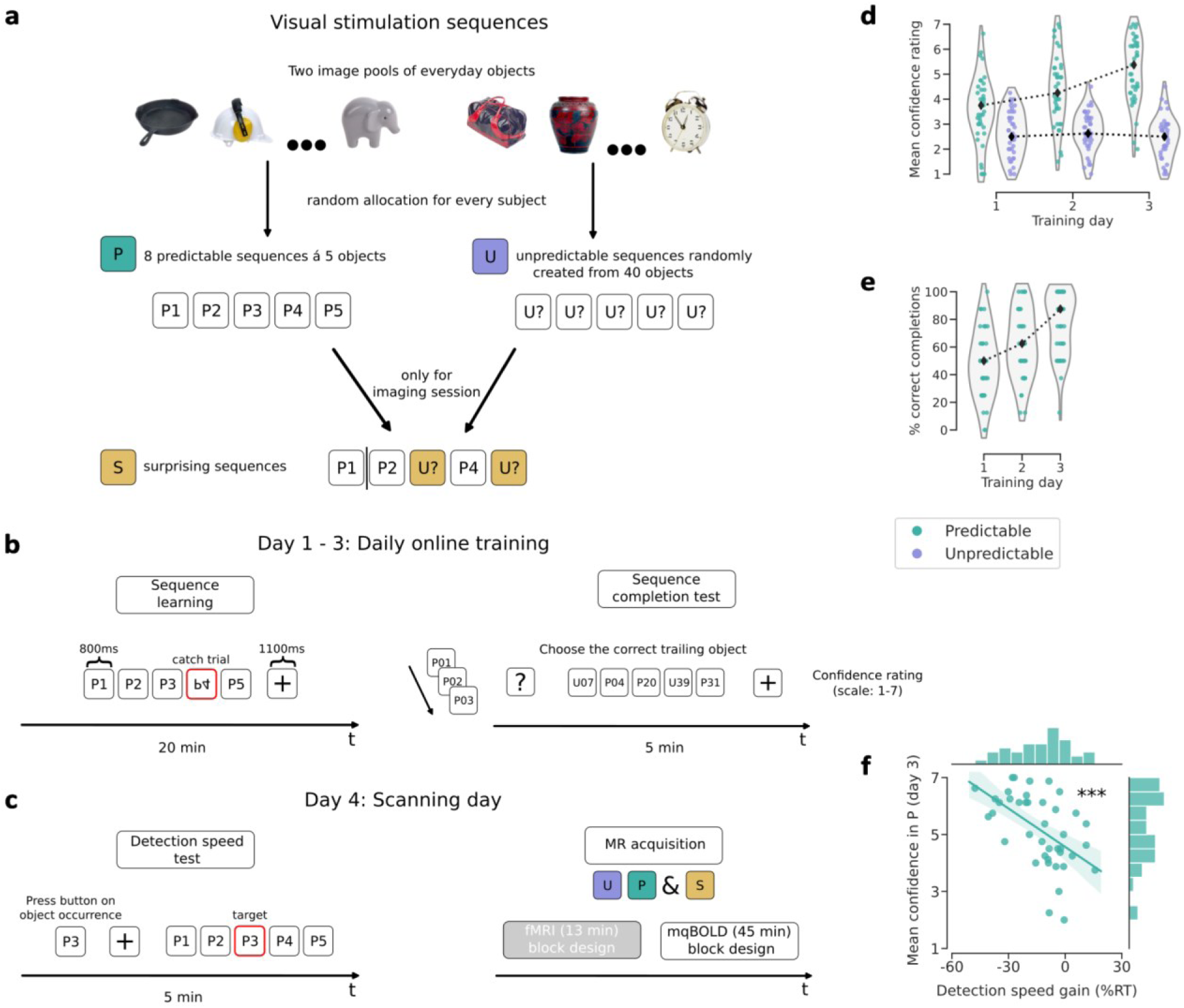
Design and behavioral results. **a**, Sequences were comprised of five everyday objects each. Predictable sequences (P) had consistent order and composition, unpredictable ones (U) were random. P and U were combined to form surprising sequences (S) that were only shown during brain imaging (Methods). **b**, Learning sessions: For three days, participants were presented with P and U sequences. Independent of predictability, participants had to react to upside-down objects. After each day, we tracked the learning progress by testing accuracy and confidence when choosing the follow-up object to incomplete sequences. **c**, Procedure during the fourth day: Differences in visual processing were measured by comparing reaction times to target objects in predictable versus unpredictable sequences. Afterwards, we acquired both energy metabolic imaging data and conventional fMRI data. See Figure 2 and Methods for further details. **d**, Mean confidence ratings for sequence completion after respective learning sessions. **e**, Mean accuracy for sequence completion after respective learning sessions. Unpredictable sequences were random and had no correct trailing object. **f**, Significant correlation between differences in target detection speed and confidence after training (r=-0.54, p<0.001). Negative values on the x-axis indicate faster reactions for predictable compared to unpredictable targets.

Prior to brain imaging on the fourth day, we included an additional task to gauge potential behavioral improvements resulting from sequence learning (Figure 1c). To this end, we again presented predictable and unpredictable sequences. Prior to each sequence, one of its objects was shown and defined as a target. Participants were instructed to react to the occurrence of the target with a button press. We reasoned that the anticipation of upcoming objects in the predictable condition would lead to faster reactions. Indeed, participants detected targets during predictable sequences more than 10% faster than in unpredictable sequences (median RT change=-10.75%, t=-5.7, p<0.001). We also found that the level of confidence after training was a strong predictor of these behavioral effects (Figure 1f).

### Experimental design replicates fMRI results of predictive processing

During the scanning phase, we acquired conventional BOLD and novel, quantitative BOLD data. In both modalities, we used a block design where multiple sequences form the unit of analysis (Methods). We used the former to validate our experimental design, comparing it to previous studies on predictive processing of visual sequences (described in Hechler et al., 2024). Our results replicate activity increases for surprising input in the ventral visual stream and activity increases linked to general prediction in the superior parietal lobe, in line with a recent meta-study (Ficco et al., 2021; Richter et al., 2018; see Supplementary Figure 1). We thereby confirmed that our task elicits established patterns of predictive processing across the cortex.

### A model for the energetic cost of predictive processing

We then investigated the net energy consumption across the cortex during processing of predictable, unpredictable, and surprising visual sequences. Unlike conventional BOLD, mqBOLD provides a rate of energy consumption, expressed as the absolute amount of oxygen consumed per minute. This rate is termed CMR_O2_ (cerebral metabolic rate of oxygen) and can be compared across different parts of the cortex. This means that mqBOLD allows the calculation of the system- or network-level energy demand and is not limited to detecting clusters of relative signal changes (as with classical BOLD fMRI).

In the present study, we aimed to provide a holistic understanding of energetic effects during predictive processing. Intuitively, local activity decreases in sensory areas might be balanced or even opposed by distributed processing across the cortex. We therefore quantified CMR_O2_ within functional networks that cover the whole processing hierarchy. We used an established cortical atlas (Schaefer et al., 2018) that defines functionally homogeneous regions and networks, differentiating large-scale sensory from higher cognitive systems (Figure 2b). These networks have been found to be stable across individuals and are in line with previous cortical parcellations schemes (Yeo et al., 2011; for details see Methods). All cortical data were fitted using a single linear mixed model to predict the energy consumption across these networks during predictable, unpredictable and surprising visual input, moderated by subjective confidence and reaction time during catch trials. We performed stepwise model selection, sequentially adding predictors of interest (Methods and Supplementary Table 1). The winning model showed that cortical energy consumption was best explained by an interaction of predictability and confidence, with respective differences for cortical networks. The fitted model is presented in Figure 2c. All model estimates are relative to the reference categories of the respective predictors: The visual network and the predictable condition. Parameter estimates for the reference levels can be interpreted for all networks or conditions for which no significant interactions are present. In the presence of significant interactions for all other predictor levels, the reference estimate only concerns the predictable condition and visual network.

**Fig 2.**
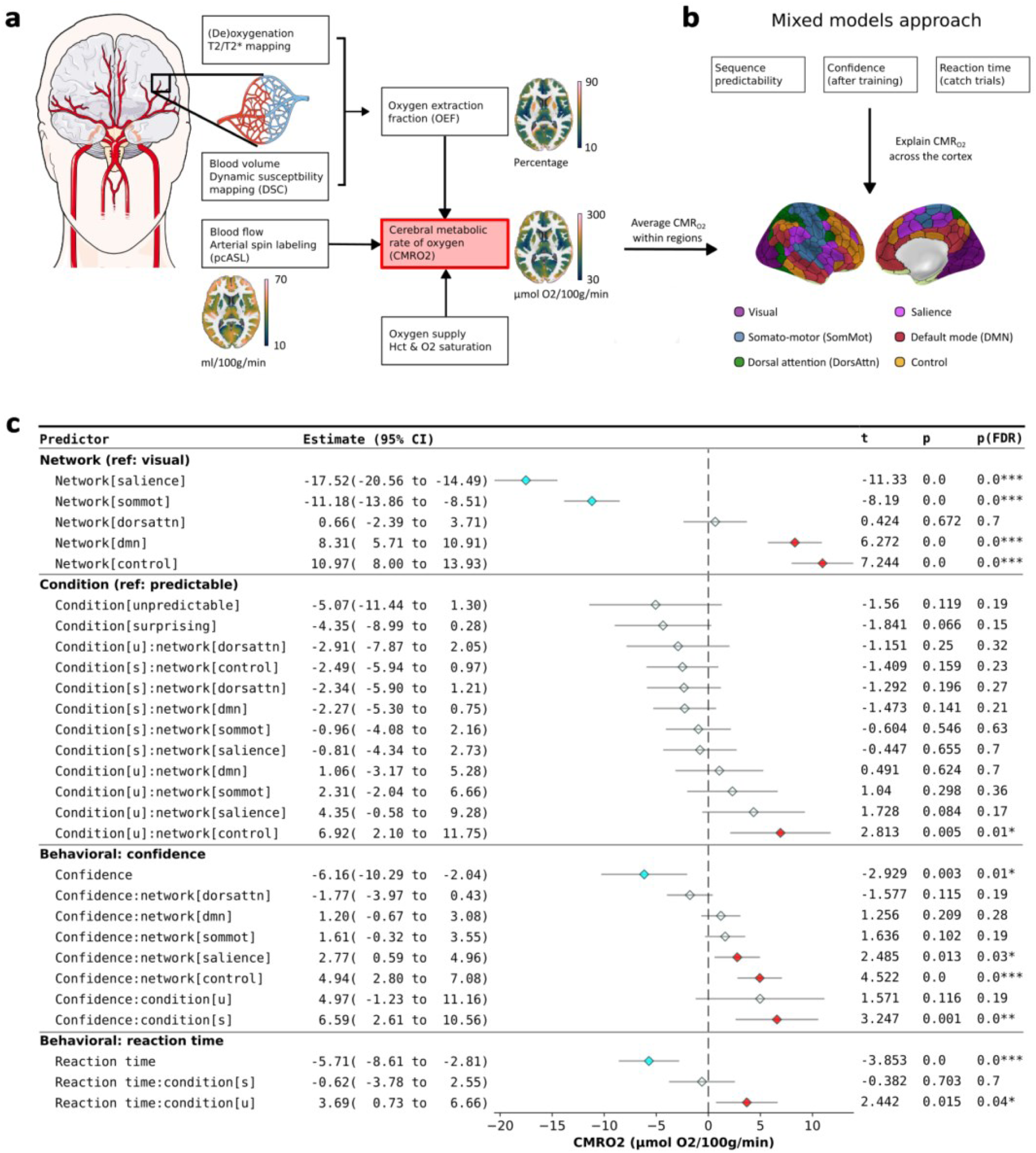
Data processing and model overview. **a**, flowchart depicting the workflow of CMRO2 calculation from experimental data*. The processing was constrained to cerebral grey matter. Parameter maps show group-level averages. Details are described in the methods section. **b**, Voxel-wise CMRO2 data was averaged within functional regions and subsequently predicted by experimental condition, confidence and reaction time using a linear mixed model. Network names and parcel locations are based on the parcellation scheme. **c**, Parameter estimates obtained from the winning model. Estimates are grouped by the main predictors and include the respective interaction terms denoted by a separating colon. Descriptions in brackets specify the level of the categorical predictors. Network abbreviations are defined in **b**, condition descriptors have been abbreviated as follows: Unpredictable = u, surprising = s. The terms within groups have been ordered by their estimate, ranging from lowest to highest. Terms surviving FDR correction have their markers colored in blue for a significant negative effect and red for a significant positive effect. *\*Visualizations of head and vessels were created using Servier Medical Art, provided by Servier, licensed under a Creative Commons Attribution 3.0 unported license*

We validated our results in two ways: As the chosen number of cortical regions in each network is arbitrary, we replicated our model with a different parcellation resolution. Furthermore, we tested whether our results were driven by outliers. To this end, we applied a robust mixed model which uses a loss function that applies less weight to extreme data points. Both alternative models yielded highly similar qualitative and quantitative results, supporting the robustness of our approach (Supplementary Figures 2 to 4).

### Confident predictions promote energy efficiency

Our model is unique in its neurobiological interpretability, as model parameters represent estimated (changes in) oxygen consumption, in units of μmol O_2_/100g/minute. Of note, oxygen is the main metabolic resource of ATP synthesis, meaning that a higher consumption indicates increased energy usage. While the model intercept is not usually interpreted, here it represents the baseline energy consumption across subjects. This baseline refers to the visual network during predictable input, for a hypothetical subject with average confidence and reaction time. We obtained a value of 138.4, which is well within the range of 120-160 usually reported for cerebral CMR_O2_ in healthy human subjects (summarized in Xu et al., 2009, see also Christen et al., 2012; Göttler et al., 2019). All estimates shown in Figure 2c are predicted deviations from this baseline (the dashed line), given a change in network, condition, confidence or reaction time.

The first group of predictors (Network, Fig. 2C) shows how networks vary in their energy consumption. Controlling for the other variables, higher cognitive networks (control and default mode) are the costliest. This is in line with recent work showing that the DMN and control network maintain highly costly connections across the cortex (Castrillon et al., 2023). The somato-motor and salience networks have the lowest rate of energy usage while the dorsal attention and visual networks occupy a middle ground.

The second group of predictors (Condition, Fig. 2C) indicates how cortical costs change with the predictability of visual input. While including this predictor and its interactions significantly improved model fit, only one of the individual terms is significant: The control network shows increased energy consumption during unpredictable input. The absence of significant estimates for the surprising and unpredictable condition suggests that input predictability alone does not trigger further detectable CMR_O2_ differences.

The third predictor group (Confidence, Fig. 2C) reveals that these effects emerge when considering predictability and confidence together. The first term shows that CMR_O2_ decreases with increasing confidence during predictable input. This means that more confident subjects use less energy on processing predictable visual input than less confident subjects, even though the stimuli were closely matched. This effect was distributed throughout the cortex, affecting sensory networks (visual, somato-motor), higher cognitive networks (DMN) and the dorsal attention network. While the interaction effects with the control and salience network are opposite in direction, it is important to note that the reference effect is negative. We therefore used simple slope analyses (Methods) to test whether the combined effect of reference and interaction estimate is significantly different from zero (Figure 3b, left). We found no evidence that confidence influences CMR_O2_ in the control (estimate=-1.23, t=-0.58, p=0.57) or salience network (estimate=-3.39, t=-1.58, p=0.12) during predictable input. Summarizing these findings, we found that the energy-metabolic benefits of confident predictions extend beyond local effects, affecting large parts of the cortex.

**Fig 3.**
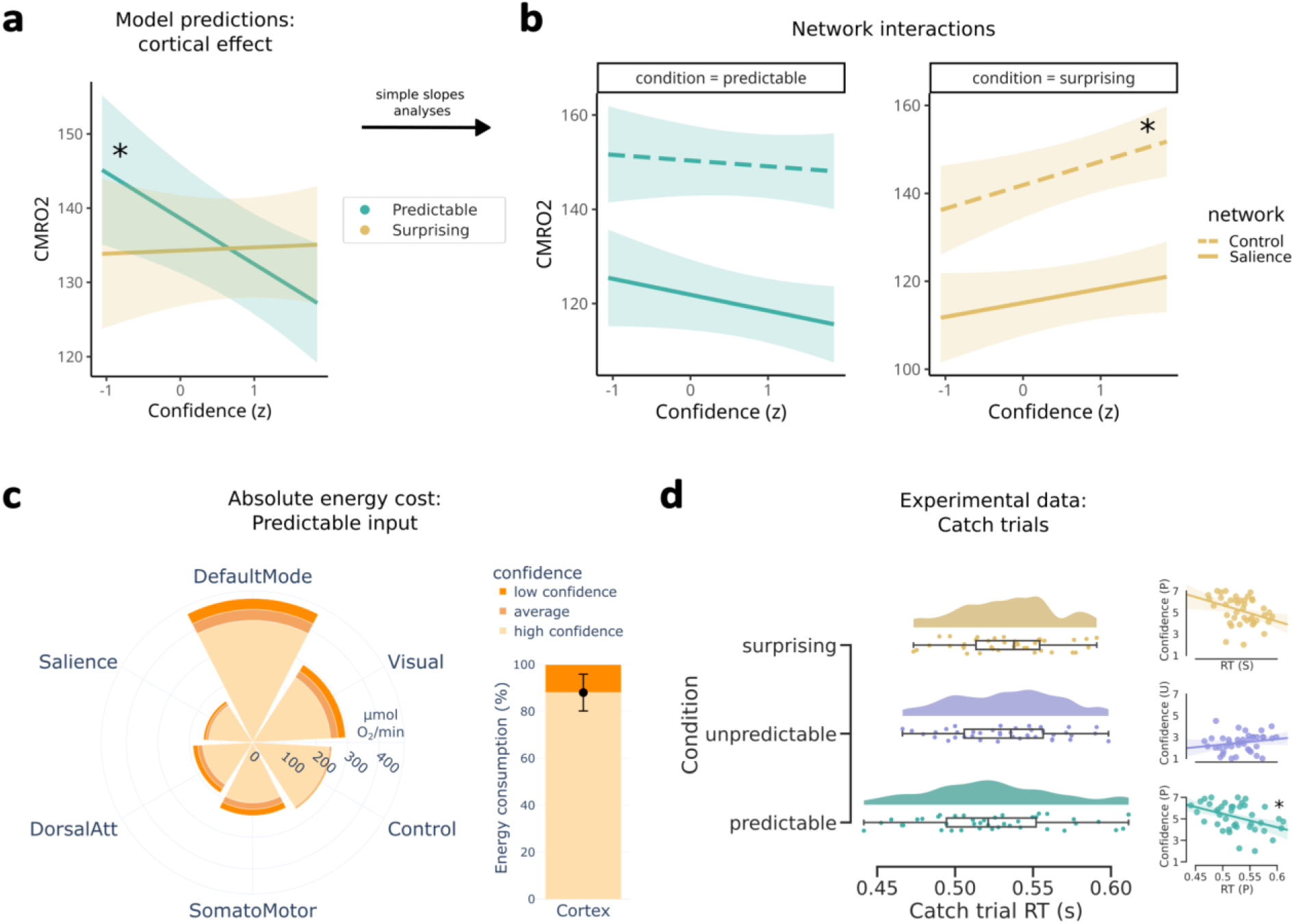
Model predictions on the cost of predictive processing. **a**, Estimated slopes for the interaction of confidence and predictability. The slope for surprising input is the linear combination of reference effect (predictable) and interaction term as determined by a simple slope analysis. **b**, Simple slope analyses for networks showing a significant interaction with confidence. In both plots, dashed lines refer to the control network. Only the positive slope of the control network during surprising input was significant. **c**, Network-wise absolute energy consumption for three levels of confidence. The bar plot shows the cortical energy consumption of highly confident subjects relative to weakly confident subjects, based on the sum of the network-wise estimates seen in the polar bar plot. **d**, Subject-wise averages of reaction time for catch trials during scanning. The scatterplots show the respective correlation with confidence ratings after training.

Regarding the link between confidence and surprising input, we found that the effect is in the opposite direction (Confidence, last row, Fig 2c). As before, we ran a simple slopes analysis, first on the reference effect. This yielded no evidence of a change in energy consumption with confidence during surprising sequences (estimate=0.42, t=0.21, p=0.84). In the context of Bayesian inference, this is an unexpected result: With increasing confidence, deviations should become increasingly impactful (being less and less in line with predictions). We therefore conducted an additional simple slope analysis for the two networks that showed no energy-saving effect of confidence during predictable input (control and salience, Figure 3b, right). This revealed that the control network consumes more oxygen during surprising input when confidence increased (estimate=5.36, t=2.57, p=0.01) and no such an effect in the salience network (estimate=3.2, t=1.5, p=0.13).

Summarizing, we found that the energy cost of visual perception is modulated by the interaction of stimulus predictability and subjective confidence. More specifically, confident participants consumed less oxygen across the cortex when presented with predictable visual sequences. On the other hand, the control network showed unique increases in energy demand during surprising and unpredictable sequences. Taken together, the energy metabolic effects of predictive processing are highly distributed throughout the cortex. Confidence is a major modulator of the resulting cost, with effects on the processing of surprising input being more spatially constrained.

Since we found widespread reductions in energy consumption for predictable input, we quantified the amount of oxygen that is saved at high levels of confidence. To this end, we used our model terms to predict CMR_O2_ for low, medium, and high confidence (based on the 5th, 50th, and 95th percentile of standardized confidence) in each network during predictable input. To account for the uncertainty in model estimates, we repeated this procedure for the lower and upper end of the CI of our confidence predictor (see Figure 2c). Figure 3c visualizes the resulting rate in μmol O_2_ per minute (scaled to network size). It can be seen that the DMN and visual network consume the bulk of the total energy budget in the cortex.

We then estimated the total energy that can be saved across the entire cortex with high confidence. We summed up the energy consumption across networks with statistically reliable effects based on our simple slope analyses: The DMN, visual network, somato-motor network and dorsal attention network. Using low confidence as a reference for maximal cost, the bar plot (Figure 3c) illustrates that high levels of confidence reduce cortical oxygen consumption by 11.96% [CI: 4.04; 19.21]. Translating back to absolute oxygen consumption, this equates to roughly 118 μmol O_2_ per minute for cortical grey matter alone (assuming an average brain weight of 1300g and a base CMR_O2_ of 140).

### Reaction times for catch trials are linked to increased energy use

The final group of predictors (Reaction time, Fig. 3c) shows the impact of reaction times across the experimental conditions. Note that these estimates concern all cortical networks. We found that the speed with which subjects reacted to the catch trials had a major impact on energy consumption, comparable to the magnitude of the confidence effect. Importantly, larger values signify slower reactions, meaning that CMR_O2_ decreased with slower reaction times. The interaction terms with predictability indicate that this relationship was weaker during unpredictable input. While our results from the training phase indicated no dependence of catch trial reactions on sequence predictability, we reevaluated this link for the scanning session. Here, reaction times differed between conditions (repeated measures ANOVA: F(2,80)=4.47, p=0.014). Reactions during predictable sequences were significantly faster than during surprising and unpredictable sequences, although the latter difference did not survive multiple comparison correction (Figure 3d; predictable vs. surprising: t(40)=-2.94, p_FWE_=0.016; predictable vs. unpredictable: t(40)=-2.24, p_FWE_=0.09).

We then asked whether the effects of confidence during predictable input can be partly explained by reaction times for catch trials. To this end, we correlated both variables to obtain the expected change in reaction time for a change in confidence (Figure 3d). We found that subjects with high confidence (associated with lower energy cost) tended to react quicker (associated with higher energy cost) when sequences were predictable (r=-0.42, p_FWE_=0.02; p_FWE_ > 0.05 for unpredictable and surprising). Consequently, reaction times offer no alternative explanation for the energy savings promoted by high confidence – rather, the energetic impact of increased reaction speed opposes this effect.

## Discussion

There is ample evidence from cognitive and computational neuroscience suggesting that we develop and optimize predictions about our environment, a process that is implemented in the brain as a hierarchy of prediction and prediction error units. However, while the behavioral advantages of accurate predictions are well documented, all functional mechanisms of the brain must operate within a limited energy budget. From an evolutionary perspective, any process improving the accuracy of perception must be weighed against the sustainability of its energetic costs. In the present study, we used an advanced metabolic imaging method to ask how predictive processing impacts energy consumption in the brain during visual processing.

We employed a sequence prediction task that addressed the two determinants of perceptual inference: The objective structure of external stimuli and the subjective accuracy of internal predictions. We manipulated the former by presenting sequences that were either deterministic, random, or surprising and sampled the latter by acquiring confidence ratings throughout a learning phase. Our results show that the brain’s energy consumption is lowest when sequences are deterministic (predictable), and predictions are precise. Importantly, this effect was found across both sensory and higher cognitive networks, amounting to a cortical reduction in energy usage of up to 12 percent. These savings did not result from reduced engagement or performance – on the contrary, predictable items were detected even quicker than unpredictable ones, a difference that also scaled with confidence. This study provides the first evidence that predictive processing is not just behaviorally but also energetically efficient.

### Energy metabolism and efficiency

When formulating biologically realistic models of brain function, accounting for resource constraints is crucial (Roberts et al., 2014; Sachdeva et al., 2021). Healthy brain function relies on constant oxygen and glucose delivery and shortages can have severe consequences ranging from cognitive deficits to cell death (P. Lee et al., 2020; Warren & Frier, 2005). Interestingly, the brain seems to be optimized for efficiency, from the firing patterns of neurons to the architecture of functional networks (Yu & Yu, 2017; Zhou et al., 2022). In related cognitive research, it has been argued that learning is an efficient process that maximizes performance while minimizing cost (Commins, 2018). These lines of research can be unified from an information-theoretic perspective: Akin to training algorithms in machine learning, the brain might aim to optimize behavioral accuracy while minimizing the complexity of internal representations (Zénon et al., 2019). This notion has been generalized under the framework of active inference (Friston et al., 2017): Humans build an internal model of the world that is continuously updated with new information, balancing model accuracy with model complexity. Assuming that the brain represents this model, the efficiency of the model might extend to the energetic efficiency of the underlying neural activity (Sengupta et al., 2013). Computational studies using neural networks support this hypothesis: Networks trained to minimize their activity in a sequence prediction task developed predictive architectures (Ali et al., 2022) and predictive learning algorithms can be expressed as energy-minimization algorithms (Luczak et al., 2022). Despite these theoretical and computational underpinnings, no direct evidence is available that predictive processing leads to metabolic efficiency.

### The role of subjective confidence during sequence learning

Studies on the human ability to extract transitional probabilities from sequential stimulation have a rich tradition in cognitive research (Schapiro & Turk-Browne, 2015). Previous work often focused on the assumed automaticity of the learning process, which can happen in the absence of intention or awareness (Alamia et al., 2016; Turk-Browne et al., 2005). However, recent work showed that, while implicit processes are a prerequisite, explicit knowledge can be acquired in parallel (Batterink et al., 2015; Dale et al., 2012). Blurring the lines further, Conway (2020) argued that most designs studying the learning process of transitional probabilities tap into the same underlying process. However, there is evidence for a difference in the neuroanatomical substrates of implicit and explicit statistical learning, so comparisons between studies should be drawn with care (Aizenstein, 2004). Recently, it has been suggested that a general Bayesian inference process underscores all probabilistic computation (Fiser & Lengyel, 2022). Consequently, instead of addressing a specific paradigm, the current work used streams of visual sequences as a tool to elicit sequence learning.

By including confidence ratings, we addressed a major parameter of Bayesian inference: Uncertainty (often referred to by its inverse, *precision*, and illustrated by the width of the prior distribution). Importantly, uncertainty has been found to govern the balance of accuracy and complexity in human inference (Tavoni et al., 2022). While Bayesian inference is a powerful model of human cognition, individuals often deviate from the idealized performance (Acerbi et al., 2014; Beck et al., 2012). This suggests that considerable interindividual differences are to be expected when submitting a group of participants to the same task. To account for this variance, we used confidence ratings as a proxy for individual uncertainty. This is in line with formulations of confidence as the posterior probability that a choice (overt or covert) is subjectively correct (Pouget et al., 2016). Interestingly, confidence ratings seem to be grounded in a “readout” of neural population activity (Geurts et al., 2022; Meyniel, Sigman, et al., 2015).

### The effect of subjective confidence on energetic cost

The link between confidence derived from computational models (ideal Bayesian observers) and brain activation during probabilistic learning has been investigated in previous studies using conventional BOLD fMRI. Meyniel & Dehaene (2017) found a small set of voxels in the inferior frontal gyrus where BOLD signals decrease with the confidence parameter when stimulus predictability is high. Similarly, Bounmy et al. (2023) reported a set of fronto-parietal and visual clusters where activity is negatively correlated with model-derived confidence. Interestingly, both studies report that subjective confidence ratings are highly correlated with the model parameter (see also Meyniel, Schlunegger, et al., 2015). Our metabolic imaging method expands on these findings in crucial ways: The BOLD signal is strongly driven by hemodynamic factors like blood flow. The link to the underlying metabolic activity, however, varies widely across brain regions (Devonshire et al., 2012). Furthermore, cognitive states affect the BOLD signal differently than the underlying oxygen consumption (Buxton et al., 2014; Moradi et al., 2012). Depending on the local coupling between neural activity and hemodynamics, BOLD signal changes and actual oxygen consumption may not even agree in their direction (Ances et al., 2008; Epp et al., 2023). As mqBOLD explicitly measures and accounts for this variance, our results are directly interpretable as energy metabolic cost, can be compared between subjects, and integrated across brain regions. While novel, mqBOLD is an established method that has been applied in healthy subjects (Christen et al., 2012) and patients with vascular disease (Göttler et al., 2019). Recently, the derived parameters have been validated against PET measurements (Kufer et al., 2022).

### Saving energy with confident predictions

We found that confidence is a major moderator of the metabolic cost of predictive processing across the cortical hierarchy. This relationship was most pronounced for deterministic visual input, where energy cost decreased with increasing confidence. In the context of predictive coding, this seems in line with the idea that feedforward transmission is limited to surprising input (Rao & Ballard, 1999) – of which there is less when predictable patterns have been learned (and are predicted with high precision). However, if perception is not driven by sensory evidence in these cases, it must be driven by prior knowledge. Predictable information generally facilitates perception (as evidenced by our behavioral findings, see also de Lange et al. (2018)), suggesting that the same, or more, information is available to the brain than sensory input alone provides. The anatomical implementation of predictive coding rests on separate prediction and prediction-error encoding neuronal populations (Bastos et al., 2012). Consequently, neural activity at each level of the hierarchy is needed during perception, irrespective of the weighting of prior versus evidence. If local reductions in activity are modulated by upstream regions, why would perception driven by prediction be more energetically efficient across cortical levels?

While our data alone cannot answer this question, we here suggest three potentially testable theories: First, the neural representation of predictable information might be more efficient. A simple example is chunking (Cowan, 2012), a psychological phenomenon where stimuli that occur together in space or time are held in working memory as a single entity. (Brady et al., 2009) specifically argued that this includes statistical regularities. With respect to our experimental design, this would suggest that confident participants represented predictable sequences as a single chunk. Related neuroimaging evidence comes from studies showing that stimuli co-occurring in time or space converge in their representation (Pudhiyidath et al., 2022; Schapiro et al., 2012).

Second, the optimization of the internal model might become more efficient. According to classical predictive coding, beliefs need to be iteratively optimized for every percept. Addressing concerns about the inefficiency of this process (regarding both time and neural activity), hybrid predictive coding has recently been proposed (Tscshantz et al., 2023).

The authors suggest that the brain learns a direct mapping of highly familiar input to the appropriate beliefs, providing an efficient “initial guess” without expensive optimization. This builds on the concept of amortized inference (Gershman, 2014), where resources are invested to develop memory resources that can be used to make future inferences less computationally intensive. The rationale here is that spending these additional resources pays off in the long run if the environment and challenges stay sufficiently stable. Regarding our design, predictable sequences presented highly stable input, making the reuse of previously learned beliefs highly effective.

Lastly, the explanation might lie in the balance of neural excitation and inhibition linked to prediction and error activity respectively. It is assumed that prediction errors are excitatory, travelling up the cortical hierarchy, while predictions are inhibitory, suppressing errors on the level below (Bastos et al., 2012; Shipp, 2016). This means that the perception of expected stimuli would minimize bottom-up excitatory activity, making inhibition more dominant. Intriguingly, inhibition is metabolically cheaper than excitation, a finding reported for both humans and mice (Vazquez et al., 2018; Waldvogel et al., 2000). This might explain why perception dominated by prior knowledge is cheaper without necessarily assuming less neural activity.

Including reaction times to the catch trials in our analysis revealed that not all reductions in energy consumption result in net savings for the cortex if participants perform a concurrent task. We found that subjects reacted quicker during predictable sequences, an effect that scaled positively with confidence. This shows that highly confident subjects improved their performance in a concurrent, but unrelated task. Importantly, reaction times and confidence were inversely related to energy consumption during predictable input. A tentative interpretation is that participants shifted their resources from model updating to lower-level perception of the objects’ orientations. Under this assumption, the brain does not aim to decrease energy consumption per se, but rather reassigns resources dynamically under the constraints of energy availability (Christie & Schrater, 2015).

Interestingly, the effect of confidence on cortical cost was not significantly different between predictable and unpredictable (random) input. Note, however, that estimates were very dissimilar on a quantitative level. Nevertheless, since we gave no feedback during the sequence prediction task, it is possible that participants thought they detected a pattern in the random sequences. This is supported by the low, but far from minimal, confidence ratings for unpredictable sequences during training. Humans tend to perceive structure even in random sequences (Huettel et al., 2002) which might drive perceptual inference in the absence of feedback. Future studies are needed to examine to which extent the effect of precise priors is independent of their objective accuracy.

### Energetic changes on a cortical level

Our study shows that an experimental manipulation as seemingly small as a change to transitional probabilities affects energy consumption across multiple sensory and higher cognitive networks. Supporting these results, the view of the brain as a collection of mostly independent modules has shifted towards accounts of a highly integrated system (Pessoa, 2023). It has recently been argued that previous approaches focusing on anatomical localization using fMRI BOLD limit our understanding of the brain to the “tip of the iceberg” (Noble et al., 2022, 2024). Specific to predictive processing, a recent study in mice found that prior expectations are represented throughout the cortex (Ishizu et al., 2024). In line with these works, we propose that the efficiency of a highly integrative system is best evaluated over a correspondingly large spatial extent.

Only the control network (also referred to as fronto-parietal network) consistently deviated from the other cortical networks in its energy usage. Here, CMR_O2_ was generally elevated during unpredictable sequences and, instead of the energy-saving effect of confidence during predictable sequences, metabolic activity increased with confidence during surprising input. The control network spans prefrontal regions (dorsal and lateral) and the posterior parietal cortex and is tightly linked to executive function and working memory (Assem et al., 2020; Menon & D’Esposito, 2022). It has been explicitly linked to processing surprising input during an attentional cueing paradigm (L. Wang et al., 2010). This might explain the increased metabolic activity when confident predictions are violated. Furthermore, the control network acts in concert with the salience network (Chand & Dhamala, 2017), especially via the anterior insula and dorsal anterior cingulate cortex (ACC). The ACC and lateral prefrontal cortex are crucial regions in a recent model of hierarchical predictive coding (Alexander & Brown, 2018). Consequently, one might have expected that the energy metabolism in the salience network is similarly sensitive to error processing. While we didn’t resolve our data on the level of specific regions, it is possible that metabolic effects only emerge when differentiating areas within this network.

### Oxidative versus glycolytically driven energy metabolism

Finally, a recent paper developed an integrative framework drawing a direct link between predictive processing and differential patterns of BOLD, CMR_O2_ and CMR_Glc_ (the cerebral metabolic rate of glucose) (Theriault et al., 2023). The authors argue that the ratio between ATP-yielding metabolites differs between bottom-up prediction errors and top-down predictions: Prediction errors rely on fast and flexible ATP generation via non-oxidative glycolysis, while prediction is based on the more efficient but less flexible oxidative phosphorylation. Importantly, the BOLD signal is driven by blood flow, which indicates oxygen supply but not necessarily oxygen consumption (Fox et al., 1988). As blood flow is more closely related to glucose use than oxygen use, BOLD is a better reflection of CMR_Glc_ than CMR_O2_ (Raichle & Mintun, 2006). It follows that, on the one hand, our CMR_O2_ data might be less sensitive to changes in error signaling but on the other hand, it could unveil prediction-related changes that BOLD does not capture. Future studies could dig deeper into these assumptions with the aim of providing a complete picture of the energy metabolism of the brain during predictive processing.

### A new approach to evaluate cognitive mechanisms

Our study presents a first step in evaluating models of cognition and perception from a new perspective: Their implementation in a biological system with limited access to energy. While quicker or more accurate sensation can improve survivability, the necessary energy consumption might not be sustainable. Intriguingly, we found that behavioral improvements are not necessarily at odds with energetic efficiency: Given high levels of confidence, predictive processing promotes fast, accurate perception while reducing processing costs. Metabolic imaging provides the unique opportunity to quantify these effects for highly integrated, distributed processing across the cortex.

## Methods

### Participants

We recruited 44 participants including students, doctoral researchers, clinic staff and the general population of Munich. All participants took part in a familiarization MRI session (20 minutes), an online training phase over three days and the main MRI session (70 minutes) on the day after training completion. Two participants were excluded due to technical problems during data acquisition. One further subject was excluded from analysis because the training phase was not properly performed. The remaining 41 participants (18 female, age [mean(std)] = 27(3.9)) were included for all analyses. The study was approved by the ethics board of the Technical University of Munich (TUM), and we acquired written informed consent from all participants.

### Visual stimuli

We selected 224 full-color images of everyday objects from a larger image set (Brady et al., 2008). The stimuli were chosen to be maximally homogenous regarding salience, e.g., by excluding food items and bright colors. All allocations of stimuli to subjects and conditions were random. Figure 1a visualizes the creation of the visual streams. A pool of 80 images was generated for every subject, with half assigned to the predictable condition and the other half to the unpredictable condition. Based on these images, eight predictable sequences were created for every subject. These objects could only occur in the sequence and position determined during stimulus creation. For the unpredictable condition, a starting set of eight sequences was created. After all were presented during the training or scanning phase, eight completely new sequences were randomly created for every repetition. Consequently, these objects never occurred in the same sequence or position, but the total number of occurrences was the same as for predictable objects. Lastly, only for the scanning phase, half of the predictable sequences and half of the unpredictable objects were combined into surprising sequences. Predictable sequences formed the basis but had one to three objects between position two and five replaced with a random object from the unpredictable condition. The first object was always unchanged to trigger conditional predictions based on the learned transition probabilities.

### Experimental design

#### Implementation

We used Psychopy (Peirce et al., 2019) to implement the design. The online training sessions were realized using pavlovia.org, where Javascript-translated Psychopy experiments can be run online with millisecond precision (Bridges et al., 2020; Sauter et al., 2020).

#### Main task

We presented participants with continuous visual streams based on the described object sequences. Each stimulus was presented for 800ms with no inter-stimulus interval within sequence and a 1100ms fixation cross between sequences. When all unique sequences of a condition were shown, their order was reshuffled for continued presentation with the constraint that objects (or sequences) could not appear twice in a row. This was done to minimize confounding effects of repetition suppression. To ensure fixation and concentration, participants were instructed to quickly react to occurrences of upside-down objects. These appeared with a probability of 10%, independent of condition.

#### Training phase

Over the course of three days prior to the main scanning session, participants followed an online implementation of the main design. This phase only included the predictable und unpredictable sequences. We instructed every participant to perform the training in a quiet environment without distractions. Stimulation blocks lasted 20 minutes per day, with a break of one minute after 10 minutes. Instructions were shown on screen before the stimulation began and the first day included a five-minute familiarization block with on-screen feedback regarding cover-task button presses. The instructions stressed the importance of the cover task, but also noted that questions regarding the order of objects in a sequence would follow each training day. During the latter task, participants saw eight incomplete sequences (the first one to four objects were shown) for both conditions (to keep image familiarity the same across conditions). After every sequence, participants chose what they assumed would be the correct trailing object from five options and gave a confidence rating on a scale of one to seven. No feedback was given regarding performance and no information on the underlying conditions was disclosed.

#### Imaging phase

During scanning, stimuli were presented against a grey background and subtended 4° of visual angle. Each condition was presented for three long blocks, during the complete duration of a pcASL sequence (6 minutes), a T2* sequence (5.5 minutes) and a DSC sequence (2.5 minutes). The resulting CMR_O2_ values were calculated as described below (see *MRI data processing*). The order was randomized, although a condition could not occur three times in a row. We left breaks of one minute between each consecutive scanning sequence. During the T2 sequence, there was no experimental stimulation. Per condition, a total of 169 object sequences over a stimulation time of approximately 14 minutes were presented. Before every scanning sequence, a reminder regarding the cover task was shown on screen. Additionally, participants got feedback on their mean reaction time after each scanning sequence to promote attentiveness and motivation. We disclosed no information regarding the object sequence patterns and the imaging session included no sequence completion test. *Object detection task*. On the day of the imaging session, prior to entering the scanner room, participants performed an object detection task on a laptop. The presentation format followed the specifications of the main task, except for transitions between sequences. We presented eight unpredictable (reshuffled) and eight predictable (as learnt) sequences in random order.

Prior to every sequence, a target object from the upcoming sequence was shown. An onscreen prompt instructed participants to react as quickly as possible to the presentation of the target image by pressing the Arrow Up button. During the task, the target image was presented for five seconds. To allow for anticipation, it could only match positions two to five of the following sequence.

### MRI acquisition

For CMR_O2_ mapping, the following sequences were acquired:

- Multi-echo spin-echo T2 mapping: 3D gradient spin echo (GRASE) readout as described previously (Kaczmarz et al., 2020), 8 echoes, TE1 = ΔTE = 16ms, TR=251ms, α=90°, voxel size 2x2x3.3mm^3^, 35 slices. T2 data was acquired once per subject, without any task.
- Multi-echo gradient-echo T2* mapping: As described previously (Hirsch et al., 2014; Kaczmarz et al., 2020), 12 echoes, TE1 = ΔTE = 5ms, TR=2229ms, α=30°, voxel size 2x2x3mm^3^, gap 0.3mm, 35 slices. T2* was acquired for all conditions.
- Dynamic susceptibility imaging (DSC): As described previously (Hedderich et al., 2019). Injection of gadolinium-based contrast agent as a bolus after 5 dynamic scans, 0.1ml/kg (maximum: 8ml per injection, 16ml per session), flow rate: 4ml/s, plus 25ml NaCl. Single-shot GRE-EPI, EPI factor 49, 80 dynamic scans, TR = 2.0s, α=60°, acquisition voxel size 2x2x3.5mm^3^, 35 slices. To stay within the limits of a full clinical dosage (16ml), we acquired DSC in two conditions only: Predictable and unpredictable. Processing of the data in the surprising condition used DSC from the predictable condition.
- Pseudo-continuous arterial spin labeling (pcASL): As described previously (Alsop et al., 2015), and implemented according to (Göttler et al., 2019; Kaczmarz et al., 2020). PLD 1800ms, label duration 1800ms, 4 background suppression pulses, 2D EPI readout, TE=11ms, TR=4500ms, α=90°, 20 slices, EPI factor 29, acquisition voxel size 3.28x3.5x6.0mm^3^, gap 0.6mm, 30 dynamic scans including a proton density weighted M0 scan. ASL was acquired for all conditions.

Prior to data acquisition, a venous catheter was placed by a medical doctor through which blood samples were taken and sent to our in-house clinical chemistry laboratory. Creatinin values were analyzed as an indicator of healthy kidney function and contrast agent was only applied for subjects below a threshold of 1.3. No subject exceeded this value. Hemoglobin and hematocrit were requested and used in modelling of CMR_O2_. Finally, arterial oxygen saturation was measured via a pulse oximeter (Nonin 7500FO, Nonin Medical B.V., The Netherlands).

### MRI data processing

To calculate CMR_O2_, the following parameters were integrated and derived as described below: The oxygen content of blood (O_2_ saturation and Hematocrit), the flow of blood (CBF), and the relative oxygen extraction (OEF). Note that the resulting CMR_O2_ values represent a consumption rate for a given condition and are not time-resolved. The processing of the quantitative parameter maps was performed with in-house scripts in MATLAB and SPM12 (Wellcome Trust Centre for Neuroimaging, UCL, London, UK). T2* images were corrected for macroscopic magnetic background gradients with a standard sinc-Gauss excitation pulse (Baudrexel et al., 2009; Hirsch & Preibisch, 2013). Motion correction was performed using redundant acquisitions of k-space center (Nöth et al., 2014). R2’ maps are derived from T2 and T2* images and yield the transverse, reversible relaxation rate that is dependent on the vascular dHb content (Blockley et al., 2013, 2015; Bright et al., 2019). However, confounds from uncorrectable strong magnetic field inhomogeneities at air-tissue boundaries, iron deposition in deep GM structures as well as white matter structure need to be considered (Hirsch & Preibisch, 2013; Kaczmarz et al., 2020). The cerebral blood volume (CBV) was derived from DSC MRI via full integration of leakage-corrected ΔR2*-curves (Boxermann, J.L., Schmainda, K.M., Weisskoff, R.M., 2006) and normalization to a white matter value of 2.5% (Leenders et al., 1990) as described previously (Hedderich et al., 2019; Kluge et al., 2016). From R2’ and CBV parameter maps, the oxygen extraction fraction (OEF) was calculated (Christen et al., 2012; Hirsch et al., 2014; Yablonskiy & Haacke, 1994). CBF maps were calculated from pcASL data, based on average pairwise differences of motion-corrected label and control images and a proton-density weighted image.

For each subject and condition, we calculated CMR_O2_ in a voxel-wise manner by combining all parameter maps via *Fick’s principle* (Fick, 1870)

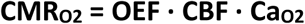

where Ca_O2_ is the oxygen carrying capacity of hemoglobin and was calculated as Ca_O2_ = 0.334 · Hct · 55.6 · O_2_sat, with O_2_sat being the oxygen saturation measured by the pulse oximeter (Bright et al., 2019; Ma et al., 2020) and Hct representing Hematocrit as measured by blood sampling prior to scanning. CBF was upscaled by 25% to account for systematic CBF underestimation due to four background-suppression pulses (Garcia et al., 2005; Mutsaerts et al., 2014). All parameter maps of each individual subject were registered to the first echo of their multi echo T2 data.

For statistical analysis of CMR_O2_, we only included voxels with a grey matter probability of > 0.5. Furthermore, the images were masked using an intersection mask to exclude voxels with excessive susceptibility, indicative of artefacts (T2 and T2* > 120ms, R2’ > 9ms) and voxels with biologically unlikely CBF (> 90 ml/min/100g) or OEF (> 90%). For each subject, we then calculated condition-wise CMR_O2_ values within 400 functional regions as defined by the Schaefer parcellation (Schaefer et al., 2018). This parcellation scheme has been used in previous influential work (e.g. Lee & Chen, 2022; Luppi et al., 2022) and was chosen for two reasons: First, regions are defined by coactivation patterns, meaning that averaging over the respective voxels upholds functional homogeneity. Secondly, it defines seven distributed functional networks that have been found to vary in their tendency to perform sensory or higher cognitive processing. These include, loosely ordered from higher cognitive to sensory (Margulies et al., 2016): Limbic network, default mode network (DMN), control network (fronto-parietal network), ventral attention network (salience network), dorsal attention network, visual network and somato-motor network. While other partitionings have been used, seven networks were shown to be both stable across subjects and compatible with previous parcellation schemes (Yeo et al., 2011). 26 of the 400 areas, mainly in the temporal pole and orbitofrontal cortex, had to be excluded from the analysis. This was due to susceptibility artefacts in the R2’ maps, leading to signal dropout in these regions. The limbic network is therefore not covered by our model.

We derived regional CMR_O2_ values in native space as follows: First, we obtained subject-wise transformation matrices from MNI152NLin2009cSym space to native space using T1w anatomical images. We registered the 400 parcel and 100 variants of the parcellation to native space, both resolving 7 networks in 2mm³ resolution. This was implemented using the *advanced normalization tools* (ANTs) toolkit via *niworkflows* in Python. Nearest neighbor interpolation was used to preserve region labels. Finally, median CMR_O2_ values over voxels in respective regions were calculated as input to our linear mixed model.

### Statistical analysis

*Mixed models*. We fitted our energy metabolic data using linear mixed models as implemented in the lme4 package in R (Bates et al., 2015). We built four models in stepwise fashion, incrementally adding predictors. We z-standardized the behavioral predictors and left the outcome CMR_O2_ values unchanged. For the complete model formulations, we refer to our code repository. The base model included fixed effects of network and reaction time to catch trials as well as random intercepts for subjects. The second model tested for an effect of the experimental manipulation by adding the condition predictor and its interactions with network and condition. The third, winning, model tested for an additional effect of confidence, including two-way interactions with network and condition. Finally, we built a model with a full three-way interaction of condition, confidence and network. Note that the within-subject design was accounted for by defining conditions as nested within subjects. The confidence ratings entering the more complex models correspond to the condition-wise average after the last training day. As surprising sequences were based on previously predictable ones, we used the confidence ratings of those sequences. Reaction times to catch trials were condition-wise averages in milliseconds. Regarding the categorical predictors of network and condition, we chose the visual network and predictable condition as reference levels. This was done to make the results more easily interpretable: Previous studies on predictive processing have contrasted predictable with surprising input to isolate prediction error processing and predictable with random input to isolate general prediction or expectation suppression effects. Model comparisons were done using likelihood ratio tests of models fitted using maximum likelihood. The reported winning model was estimated using restricted maximum likelihood. Approximate t and p-values are based on the Satterthwaite degrees of freedom method using the R package lmerTest (Kuznetsova et al., 2017). Finally, we provide corrected p-values for each predictor using the false discovery rate method.

As a follow-up, we used simple slopes analyses (using the *interactions* package in R; Long, 2022) to estimate the net energetic effect for networks that diverged from the reference effect (if both estimates were significant). Especially in the case of opposing signs, this approach can validate whether the total effect is significantly different from zero (Bauer & Curran, 2005).

To ensure that our results are not strongly driven by outliers, we validated our winning model by fitting it with a corresponding robust method as implemented in the R package robustlmer (Koller, 2016). This method uses the Huber loss function, which is quadratic for small differences, but linear for large differences.

*Model-based CMR_O2_ cost predictions*. For model-based predictions, we added the corresponding parameter estimates as described in the results. Total CMR_O2_ was therefore calculated as the sum of the intercept (the sample baseline for predictable input in the visual network), the network effect (the difference of the network energy consumption from the visual network) and the reference effect of confidence (concerning the predictable condition). We calculated the predicted cortical CMR_O2_ for weakly confident (5^th^ percentile, confidence(z)=-1.96), average (50^th^ percentile, confidence(z)=0) and highly confident (95^th^ percentile, confidence(z)=1.96) subjects. The outcome was then scaled to the grey matter mass of each network in MNI space, approximated by grey matter voxel count multiplied by a tissue mass of 0.0014 gram per cubic millimeter (Barber et al., 1970; IT’IS Foundation, 2022). Finally, predicted CMR_O2_ was summed over networks to obtain cortical values. We repeated this procedure for the 5^th^ and 95^th^ percentile of the confidence parameter estimate (Figure 2c) to obtain an upper and lower bound of the predicted energy cost.

## Data availability

All behavioral data as well as the subject-wise, region-averaged CMR_O2_ data are available as part of our github repository https://github.com/NeuroenergeticsLab/confidence_prediction_energy. These data allow the direct replication of the reported mixed model results. To protect participant privacy, the raw imaging and blood data used in the mqBOLD approach are available from the authors upon request.

## Code availability

The code to reproduce all figures and statistical analyses can be found in our GitHub repository: https://github.com/NeuroenergeticsLab/confidence_prediction_energy.

## Acknowledgements

The authors want to thank Christine Preibisch for providing MRI protocols and template code for acquisition and processing of mqBOLD MRI data. We further thank Franziska Knolle for helpful discussions on model design and interpretation.

## Author contributions

AH performed design conceptualization and implementation, investigation, formal analysis, visualization, and writing. FdL provided experimental design conceptualization and writing review. VR provided supervision, project conceptualization, funding and writing.

