## Supplementary information for "Energy savings across the cortex for confidently predicted visual input"

Contents

Supplementary figure 1 to 4

Supplementary table 1

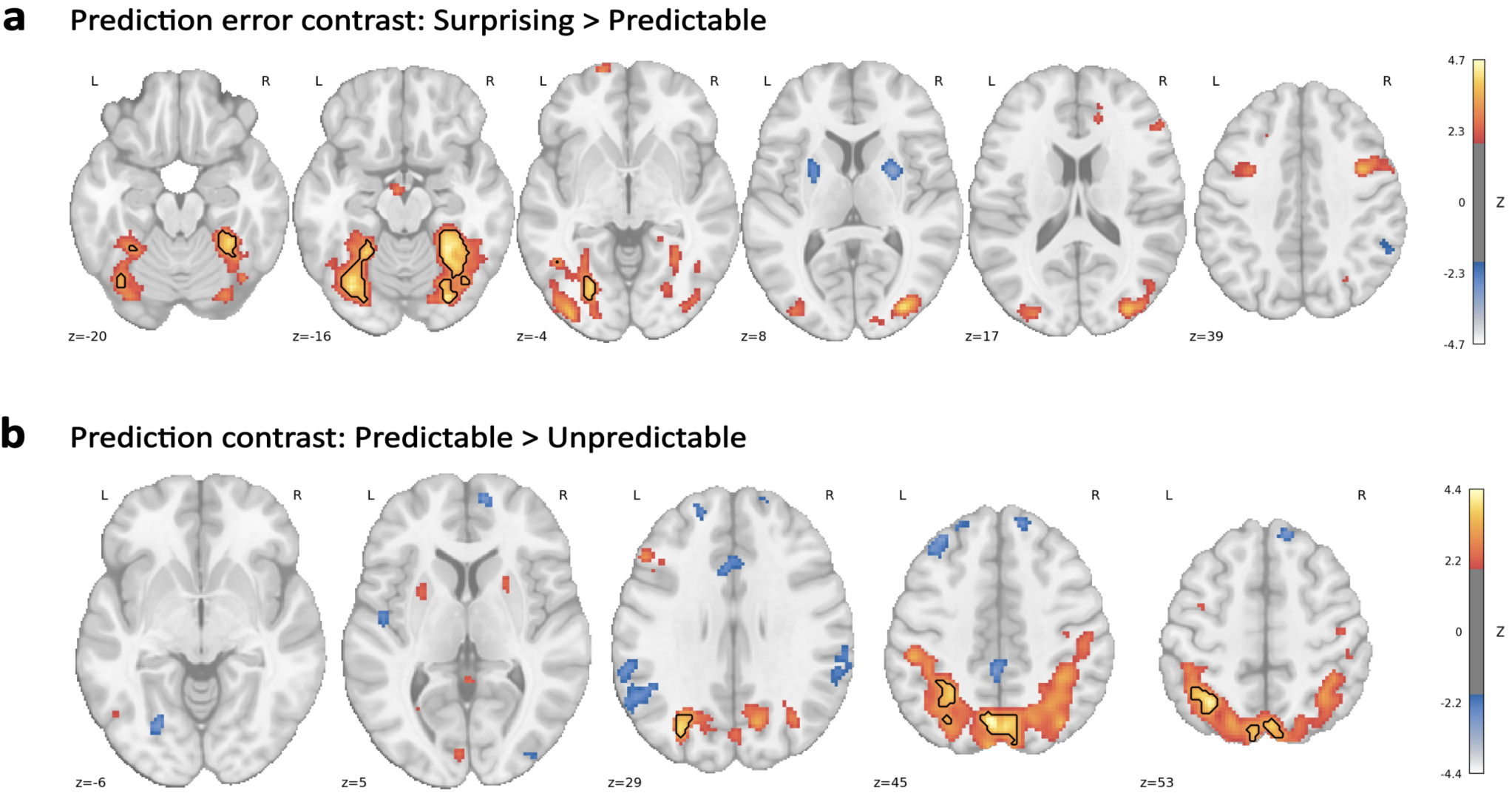

**Supplementary Figure 1 | BOLD data results.** The same participants were subjected to a standard fMRI scan before metabolic imaging, using the same experimental design. The BOLD data was analyzed using block regressors, with four sequences of one condition constituting one block. Full methodological details are given in Hechler et al. (2024). Brain slices show uncorrected t-values thresholded at z=2.33 (p<0.01). Clusters surviving cluster correction (cluster-forming threshold: z=3.09, cluster threshold: p_FWE_=0.025) are shown in black contours. **a**. Surprising > predictable contrast. **b**. Corresponding results for the predictable > unpredictable contrast.

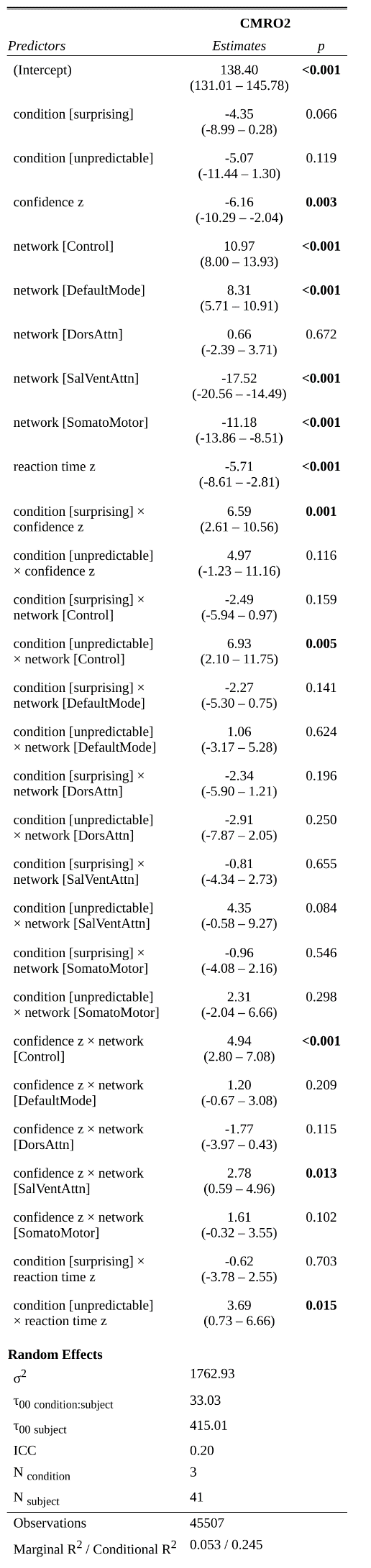

**Supplementary figure 2 | Summary of the winning mixed model.** The full formula used (following notational conventions) was:

CMRO2 ~ condition*confidence_z + network + network:condition + network:confidence_z + reaction_time_z + reaction_time_z:condition + (1|subject/condition)

Note that the salience ventral attention network (SalVentAttn) was renamed to “salience” in the main paper for better readability. We describe the details of the statistical approach in the Methods.

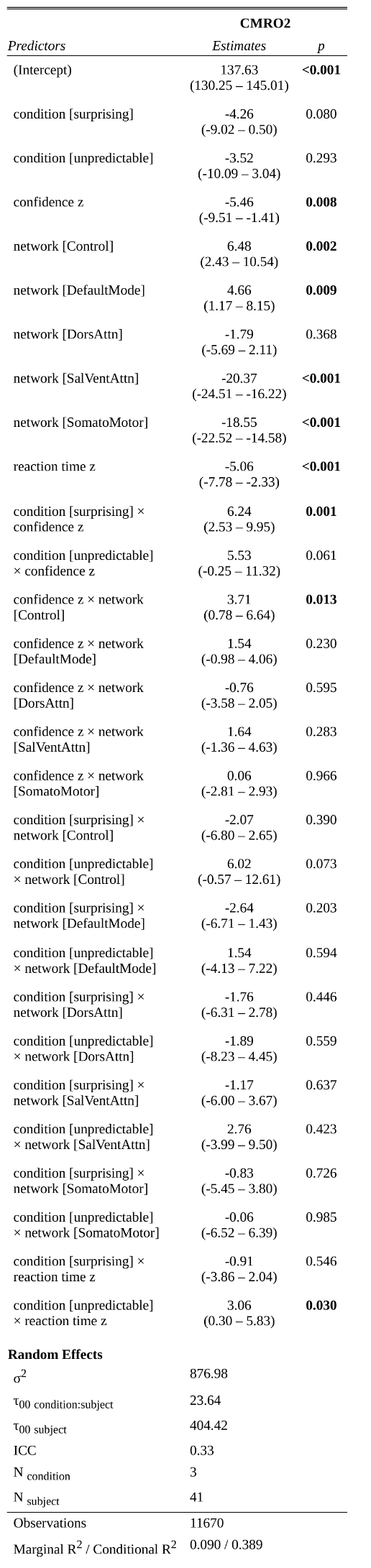

**Supplementary figure 3 |** **Summary of the replication model using 100 parcels.** We used the same formula as for the winning model. CMR_O2_ data was averaged within 100 instead of 400 parcels as defined by the Schaefer parcellation. Further details are described in the Methods.

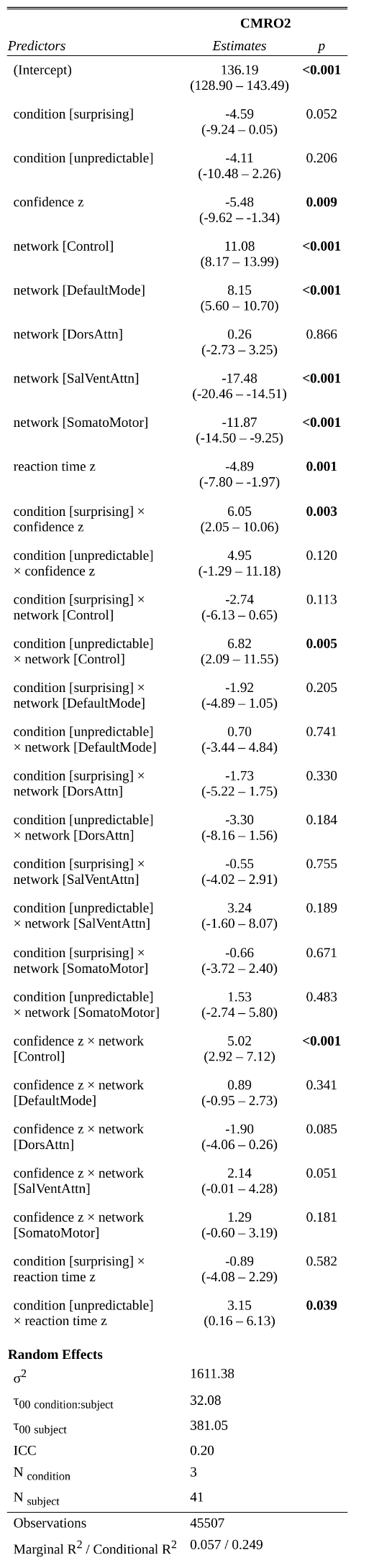

**Supplementary figure 4 |** **Summary of the replication model using 400 parcels and robust estimation.** We used the same formula and data as for the winning model. Further details are described in the Methods.

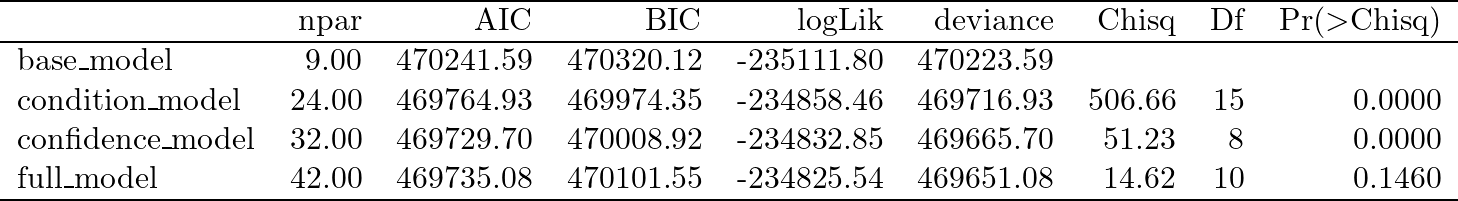

**Supplementary Table 1 | Step-wise model comparisons.** Models were sequentially tested using likelihood ratio tests. Each model includes all predictors of the previous ones. The condition and confidence models added both the respective main effects and interactions as described in the Methods. *npar*: number of parameters.
